# Hemodynamic Correlates of Fluctuations in Neuronal Excitability: A Simultaneous Paired Associative Stimulation (PAS) and functional Near Infra-Red Spectroscopy (fNIRS) Study

**DOI:** 10.1101/2021.09.29.462418

**Authors:** Zhengchen Cai, Giovanni Pellegrino, Amanda Spilkin, Edouard Delaire, Makoto Uji, Chifaou Abdallah, Jean-Marc Lina, Shirley Fecteau, Christophe Grova

## Abstract

**Background:** The relationship between task-related hemodynamic activity and brain excitability is poorly understood in humans as it is technically challenging to combine simultaneously non-invasive brain stimulation and neuroimaging modalities. Cortical excitability corresponds to the readiness to become active and as such it may be linked to metabolic demand.

**Hypotheses:** Cortical excitability and hemodynamic activity are positively linked so that increases in hemodynamic activity correspond to increases in excitability and vice-versa.

**Methods:** Fluctuations of excitability and hemodynamic activity were investigated via simultaneous Transcranial Magnetic Stimulation (TMS) and functional Near Infrared Spectroscopy (fNIRS). Sixteen healthy subjects participated in a sham-controlled, pseudorandomized, counterbalanced study with PAS (PAS10/PAS25/Sham) on the right primary motor cortex (M1). The relationship between M1 excitability (Motor Evoked Potentials, MEP) and hemodynamic responses to finger tapping reconstructed via personalized fNIRS was assessed.

**Results:** Hemodynamic activity exhibited a significant correlation with cortical excitability: increased HbO and HbR (absolute amplitude) corresponded to increased excitability and vice-versa (r=0.25; p=0.03 and r=0.16; p=0.17, respectively). The effect of PAS on excitability and hemodynamic activity showed a trend of positive correlation: correlation of MEP ratios (post-PAS/pre-PAS) with HbO and HbR ratios (r=0.19, p=0.29; r=0.18, p=0.30, respectively).

**Conclusions:** TMS-fNIRS is a suitable technique for simultaneous investigation of excitability and hemodynamic responses and indicates a relationship between these two cortical properties. PAS effect is not limited to cortical excitability but also impacts hemodynamic processes. These findings have an impact on the application of neuromodulatory interventions in patients with neuropsychiatric disorders.

## 1. Introduction

The relationship between cortical excitability and elicited hemodynamic activity is poorly understood in humans. Such investigation requires simultaneous measurements of excitability and hemodynamic activity [Siebner et al., 2009]. Transcranial Magnetic Stimulation (TMS) is a versatile technique that allows the assessment and modulation of primary motor cortex (M1) excitability. Cortical excitability assessment is easily achieved via single pulse TMS (spTMS) by measuring the amplitude of the Motor Evoked Potentials (MEPs). M1 excitability modulation can be obtained by repetitive TMS (rTMS) and related techniques, such as Paired Associative Stimulation (PAS). PAS was established about two decades ago [Stefan, 2000] and consists in combining TMS with peripheral electrical stimulation such as Median Nerve Stimulation (MNS). It relies on the principle that repetitive stimulations delivered with proper timing and pace induce long-term potentiation and long-term depression like plasticity, exploiting the concept of spike-timing-dependent plasticity [Levy and Steward, 1983; Rossini et al., 2015]. PAS effects can last for 30 minutes or more [Lee et al., 2017; Stefan, 2000; Suppa et al., 2017]. This technique is therefore well-suited to manipulate cortical excitability by tuning the timing of the interstimulus intervals (ISI), because it can generate either an excitability increase, when applying a 25ms time-interval between MNS and TMS pulses (PAS25), or an excitability decrease, when applying a 10ms time-interval between pulses (PAS10). A PAS sham has also been described and validated [Gow et al., 2004; Loo et al., 2000; Michou et al., 2014].

A standard approach to non-invasively map brain functions is to measure the fluctuations of brain hemodynamic signals elicited by neuronal activity. Functional magnetic resonance imaging (fMRI) is typically the tool of choice for its reliability, ease of use, high spatial resolution and sensitivity to deep brain regions [Bandettini et al., 1992; Glover, 2011; Kwong et al., 1992]. Nevertheless, bringing TMS within the MRI environment is challenging [Hallett et al., 2017]. MRI compatible TMS coils have been developed [Navarro De Lara et al., 2015; Wang et al., 2017] and applied to investigate the TMS induced hemodynamic responses [Navarro de Lara et al., 2017], but the majority of studies have had an “offline” approach, meaning that no continuous fMRI measurement during TMS is usually performed [Siebner et al., 2009].

Functional Near Infra-Red Spectroscopy (fNIRS) is a non-invasive neuroimaging modality, which allows monitoring changes in oxy- and deoxy-hemoglobin (i.e., HbO/HbR) in the cerebral cortex [Jöbsis, 1977; Scholkmann et al., 2014; Yücel et al., 2021]. It measures light intensity changes, modulated by local absorption associated with underlying hemoglobin concentration fluctuations, via source-detector pairs placed on the scalp. fNIRS has high temporal resolution and acceptable spatial resolution with no electromagnetic interference [Curtin et al., 2019b; Siebner et al., 2009; Vasta et al., 2017]. It allows the estimation of both HbO and HbR concentration changes in cortical regions. It is relatively portable and permits prolonged scans, with little or no discomfort for the participants [Gramigna et al., 2017; Pellegrino et al., 2016; Scholkmann et al., 2014]. The drawback of combining fNIRS and TMS is that when both techniques are applied to the same cortical area, the optodes (fNIRS sensors) introduce some additional space between the TMS coil and the scalp. This requires higher TMS stimulation intensities as the strength of the magnetic field decays sharply when increasing the coil distance to the target region [Curtin et al., 2019b; Parks, 2013]. Nevertheless, we are proposing combining PAS and fNIRS as a promising way to assess the relationship between cortical excitability and hemodynamic responses.

We conducted here the first PAS-fNIRS investigations of the relationship between cortical excitability and task-related hemodynamic response. Cortical excitability corresponds to the cortical readiness to become active and as such it may be linked to metabolic demand. We first hypothesized that fluctuations of cortical excitability are positively correlated with fluctuations of hemodynamic activity. This hypothesis was tested by estimating the correlation between MEP peak-peak amplitude and task-related HbO/HbR response, regardless of the PAS interventions (PAS25, PAS10, Sham). Second, we further hypothesized that PAS affects cortical task-related hemodynamic responses in addition to cortical excitability, and such modulations of excitability and hemodynamic positively correlate. In other words, when PAS increases cortical excitability, it results in enhanced task-related hemodynamic response, whereas when PAS decreases cortical excitability, it results in reductions of the hemodynamic response to the task. This hypothesis was tested by estimating the correlation between MEP and HbO/HbR ratios calculated as the post- over pre-intervention amplitudes.

## 2. Material and methods

### 2.1 Subjects and study design

This study was approved by the Central Committee of Research Ethics of the Minister of Health and Social Services Research Ethics Board, (CCER), Québec, Canada. All subjects signed a written informed consent before participation. Only subjects meeting the following inclusion/exclusion criteria were considered: 1) age between 18 and 40 years old; 2) right-handed male; 3) no present or past neurological disorders; 4) no medications acting on the central nervous system; 5) no contraindications to MRI or TMS [Rossi et al., 2009; Suppa et al., 2017]. Nineteen subjects (19 – 35 years old) participated in the study. Subjects were instructed to have a regular sleep cycle for the days before the experiments and to not consume caffeine for at least 90 minutes before the experiment.

The experimental design and setup are illustrated in Fig.1 and Fig.2, respectively. Every participant had 1) an anatomical head MRI scan (T1- and T2-weighted, 1mm^3^ isotropic) for neuronavigated TMS; calculate personalized fNIRS optical head model [Machado et al., 2014; Machado et al., 2018]; and install optodes, followed by 2) somatosensory evoked potentials recording during electrical stimulation of the median nerve at the wrist to measure N20 latency and tune PAS accordingly. We then conducted three experimental sessions corresponding to three different PAS interventions: PAS25, PAS10, and sham. These sessions were performed at least two days apart to minimize potential carryover effects. As this study included three sessions, we decided to consider male participants to minimize the confounding of cortical excitability changes due to the menstrual cycle [Hattemer et al., 2007; Lee et al., 2017]. Experimental sessions were performed with a pseudorandomized order, counterbalanced across subjects, which consisted of the following components:

- PAS was performed with 100 pairs of MNS and TMS with a fixed interval of 10s according to the guidelines [Suppa et al., 2017], for a total duration of 18 minutes (Fig.1c). MNS was delivered at the left wrist and with the following parameters: intensity = 300% perceptual threshold, square wave and 0.2ms duration. TMS intensity was 120% of the resting motor threshold (RMT). The ISI between MNS and TMS was then set to be individual N20+5ms for PAS25 and N20-5ms for PAS10, respectively [Carson and Kennedy, 2013]. Sham was the same as PAS25, except TMS was not delivered, whereas its sound was provided via a stereo speaker [Zangrandi et al., 2019].

Before and after each intervention,

- Cortical excitability was measured via M1 spTMS, applying 75 stimuli (ISI ranging between 5s and 25s) to the right hand-knob (Fig.1b). MEPs were recorded by bipolar electromyography (EMG) electrodes attached on the right abductor pollicis brevis (APB), with a standard belly-tendon montage (Fig.2d). The RMT was defined with the TMS Motor Threshold Assessment Tool [Ah Sen et al., 2017; Awiszus et al., 1999]. All TMS procedures followed the recommendations of the International Federation of Clinical Neurophysiology, and no participants reported any severe discomfort or side effects [Rossi et al., 2009].
- A finger tapping task was performed to activate M1 and estimate its task-related hemodynamic activity. Subjects were instructed to tap the left thumb to the other digits sequentially, at a pace of about 2Hz (Fig.1a). The movement was performed in short blocks of 10s interleaved with a resting period jittered between 30s to 60s. This time-constraint was meant to avoid task events phase locking to undergoing physiological hemodynamic oscillations [Aarabi et al., 2017]. Movement onset and offset were instructed by auditory cues. The duration of the motor task was about 18 minutes and consisted of 20 blocks.

**Fig.1.**
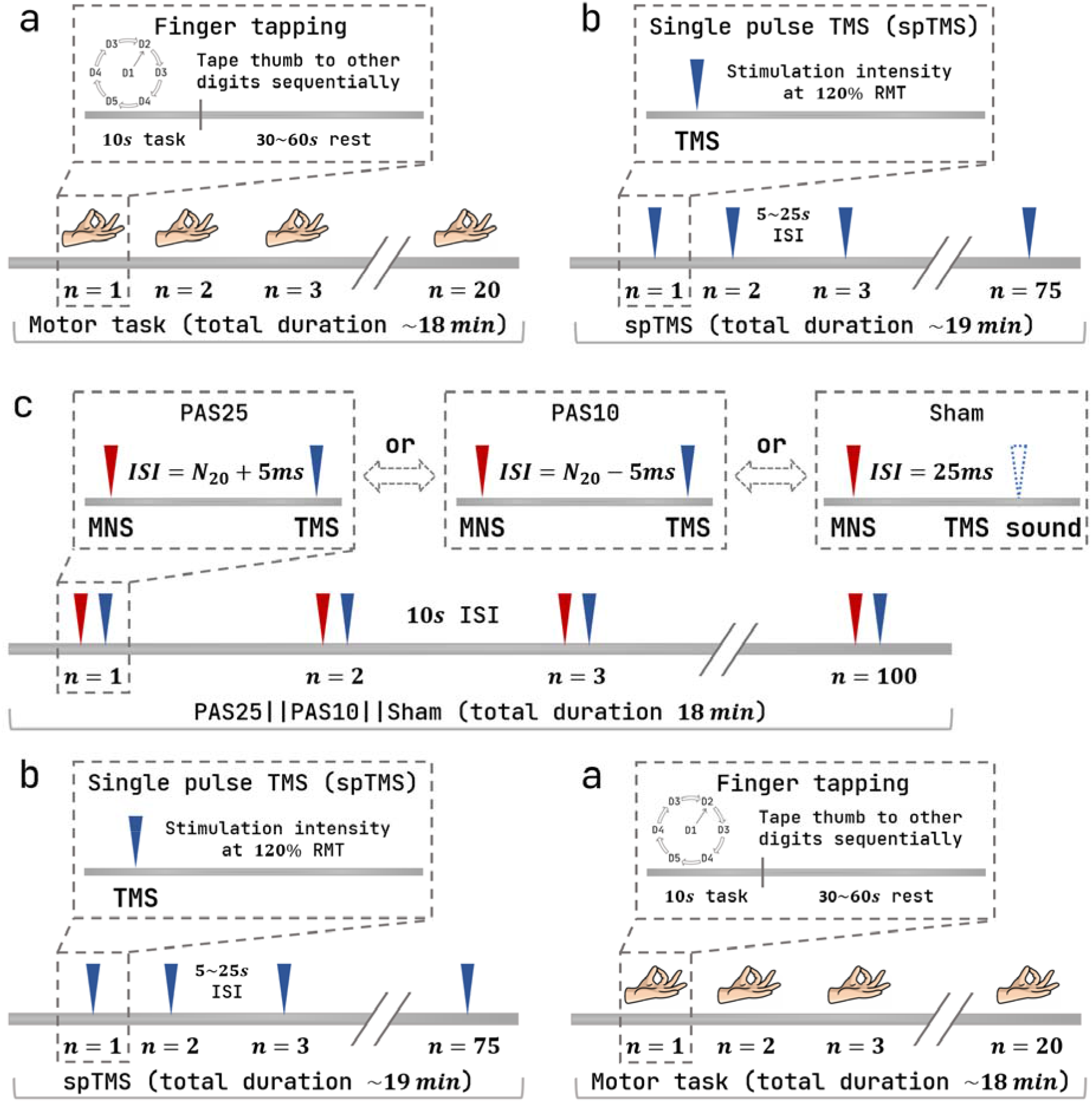
Study design. Each acquisition session described above contains sections following the order of a, b, c and b, a. (a) Finger tapping task: subjects were asked to tap the left thumb (D1) to the other four left digits (D2-D5) sequentially, at around 2Hz. Each tapping block lasted 10s and was followed by 30s to 60s jitter rest. Start and stop signals were delivered by auditory cues. 20 blocks were performed for 18 minutes total duration. (b) A single pulse TMS was delivered onto the “hot spot” at 120% of individual RMT. 75 pulses were delivered with a 5s to 25s jittered ISI for a total duration of about 19 minutes. (c) Three PAS interventions were performed on different days separated by at least 2 days and presented in a pseudorandomized order. ISI between peripheral (MNS) and TMS was set to individual N20+5ms (PAS25) and N20-5ms (PAS10) for excitatory and inhibitory PAS, respectively. Sham was similar to PAS25, same MNS followed by a 0 intensity TMS, whereas the TMS click was reproduced via speakers. MNS intensity was set to 3x individual perceptual threshold. PAS pairs were separated by a 10s interval. In total, 100 pairs were delivered in 18 minutes. Note that fNIRS signal was acquired during the whole experiment session.

**Fig.2.**
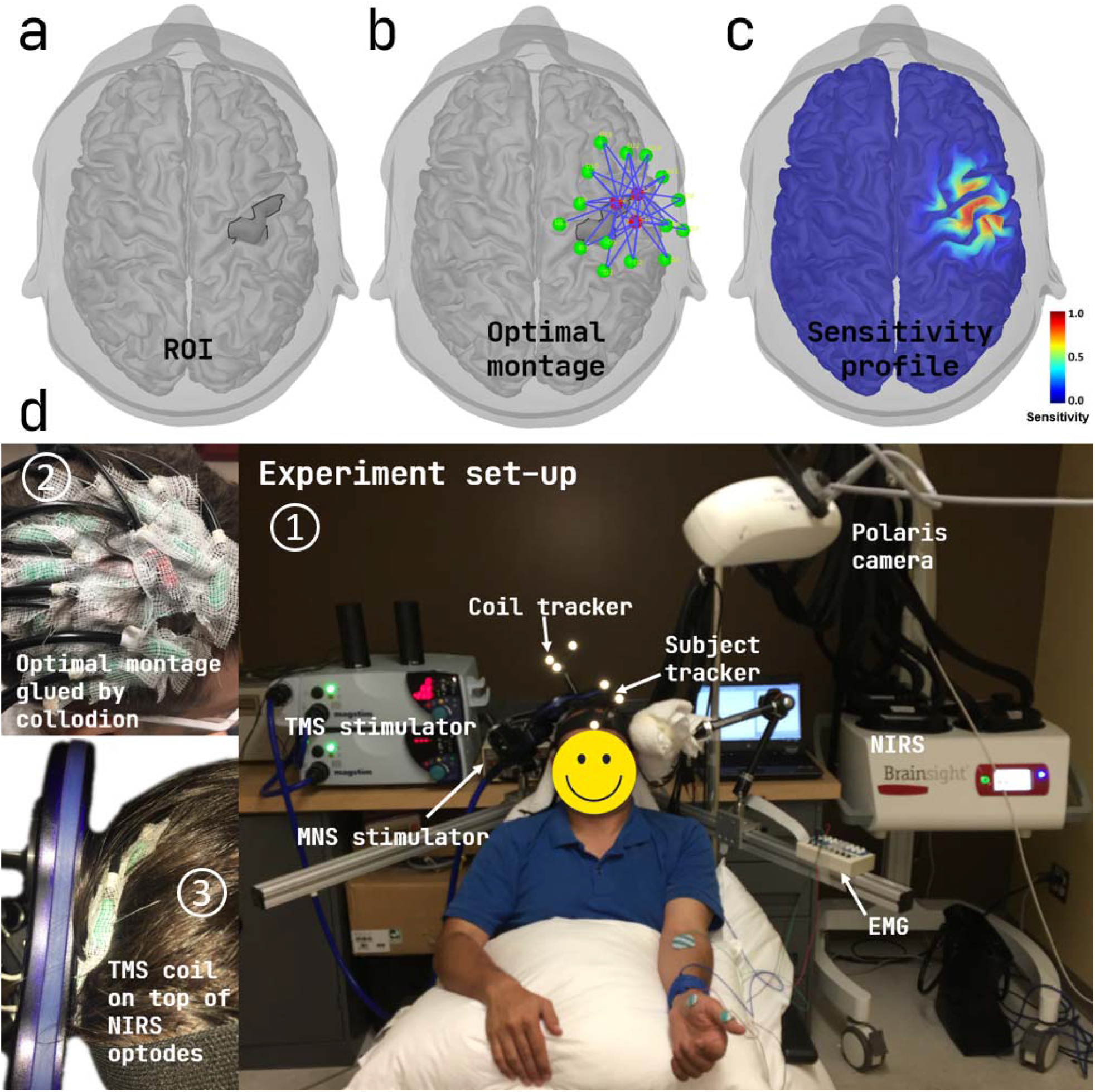
Personalized optimal montage and simultaneous PAS-fNIRS acquisition setup. (a) The right M1 hand region (“hand knob”) was pre-defined as the target ROI of personalized optimal fNIRS montage. This area was manually selected on each subject’s cortical surface extracted from individual MRI. (b) Example of the optimal montage estimated for Sub06. The montage was constrained to have 3 sources (red), 15 detectors (green) and 1 proximity detector (placed in the middle of sources, not shown). (c) Normalized light sensitivity profile of the optimal montage expressed as the sum of all channels’ sensitivity along the individual cortical surface. (d) PAS and fNIRS acquisition setup. (1) Participants sat on a comfortable TMS armchair. Four different machines were involved: a Brainsight-fNIRS for fNIRS data acquisition and neuronavigation; Magstim 200^2^ TMS stimulators for spTMS and PAS; a Digitimer DS7A for MNS and PAS; and a BrainAmp ExG bipolar system for recording MEPs. (2) Low-profile (thin) optodes were attached on the subject’s head using collodion based on their optimal positions defined in (b). (3) A figure-8 TMS coil was placed on top of the optodes, guided with the neuronavigation system, held with a mechanical arm.

fNIRS data were acquired using a Brainsight fNIRS machine (Rogue-Research Inc, Montreal Canada), sampling at 10Hz. MEPs were recorded by a BrainAmp ExG bipolar system (Brain Products GmbH, Germany). Please refer to Appendix A for further detailed experiment protocol. From the nineteen subjects, one was excluded due to low sensitivity to TMS and two due to poor fNIRS signal quality. Four subjects dropped out after the first session due to personal reasons, resulting in 16 PAS25, 12 PAS10 and 12 sham sessions.

### 2.2 Personalized fNIRS using the optimal montage

In order to maximize the sensitivity of fNIRS channel layout (i.e., montage) to M1 hemodynamic activity, we applied a personalized optimal montage, previously developed and validated by our group. Specifically, this montage maximizes fNIRS sensitivity along the cortical surface with good spatial overlap between channels [Cai et al., 2021b; Machado et al., 2014; Machado et al., 2018; Machado et al., 2021; Pellegrino et al., 2016]. The T1-/T2-weighted images were processed by FreeSurfer 6.0 to segment the head (i.e., scalp, skull, cerebrospinal fluid, gray matter and white matter) and generate a mid-cortical surface (i.e., a middle layer of the gray matter, between pia mater and gray-white matter interface) [Fischl et al., 2002]. The target area for the optimal montage (see Fig.2a) corresponded to the right hand-knob and was manually defined for each participant along the mid-surface [Raffin et al., 2015]. The personalized optimal montage was estimated imposing the following constraints: 1) 3 light sources and 15 detectors (see Fig.2b); 2) distance between source-detector pairs ranging from 2.0cm to 4.5cm and 3) large spatial overlap between channels, e.g., each source must construct at least 13 channels among 15 detectors. The output of the optimal montage procedure is a set of fNIRS optode positions along the scalp to probe the right hand-knob with the highest sensitivity. Finally, a proximity detector for recording physiological hemodynamics of the scalp was added at the center of the 3 sources [Gregg et al., 2010; Zeff et al., 2007].

### 2.3 Excitability data analysis

EMG data collected during spTMS were analyzed using the Brainstorm software [Tadel et al., 2011] and R 4.0.3 [R Core Team, 2020]. They were filtered between 3 and 2000Hz. MEP trials were extracted within a time window from -10ms to 100ms around the stimulation and baseline corrected (−10ms to 0ms). Run specific (e.g., pre-PAS 25 of Sub01) excitability was expressed as the average of MEP peak-peak amplitudes across spTMS. Session specific (e.g., PAS 25 of Sub01) excitability change was measured as the post-/pre-PAS ratio of averaged MEP peak-peak amplitudes.

### 2.4 fNIRS data processing

The details of the following procedures are provided in Appendix B. Briefly, fNIRS data processing was performed applying 3D reconstructions with the Maximum Entropy on the Mean (MEM) framework, as described in [Cai et al., 2021a; Cai et al., 2021b]. The goal was to extract HbO/HbR amplitude from spatiotemporal maps reconstructed from the channel space data to the underlying cortical surface. fNIRS data pre-processing involved the following steps: bad channel rejections; physiological noise regression using proximity channels [Gregg et al., 2010; Zeff et al., 2007]; band-pass filter between 0.01Hz and 0.1Hz, and epochs extraction with a time window of -10s to 30s around the task onset. To extract robust and reliable HbO/HbR amplitude for further estimation of PAS effects on hemodynamic, we introduced in this study an original approach which comprises three steps: 1) selection of 101 trials centred around the median signal to noise ratio (SNR) of “all the possible” sub-averaged 16 out of 20 fNIRS epochs. This procedure aimed to exclude eventual motion artifacts contaminated epochs from sub-averaging and resulted in a distribution representing the variability of task-evoked fNIRS signal changes specific for each run (e.g., pre-PAS25 of Sub01); 2) fNIRS 3D reconstructions of each sub-averaged trial using the MEM framework, resulted in 101 spatiotemporal hemodynamic responses maps for each task run; 3) extraction/identification of a data-driven ROI which exhibited significant task-related hemodynamic responses for each session (e.g., PAS25 of Sub01). Importantly, since this procedure was data-driven, fNIRS analysis was blind to PAS interventions. Finally, the measure of HbO/HbR was defined as the averaged amplitude within a 5s window centred around the peak of the reconstructed time course, extracted by averaging reconstructed time courses within the ROI defined in 3). Therefore, 101 HbO/HbR amplitudes were extracted for each run. Run specific hemodynamic responses were expressed as the averaged HbO/HbR amplitude among all 101 trials. Session specific PAS effects on task-related hemodynamic response were represented by the post-/pre-intervention ratios of averaged HbO/HbR measures. To demonstrate group-level intervention effects on reconstructed HbO/HbR maps, individual maps selected at their respective peak were first coregistered on the mid-surface of MNI ICBM152 [Fonov et al., 2009; Fonov et al., 2011] template using FreeSurfer spherical registration, and then averaged over all subjects.

### 2.5 Statistical analysis

To assess the relationship between fluctuations of cortical excitability and task-related hemodynamic activity regardless of PAS intervention, we pooled together subjects’ data of all runs (before and after PAS) and calculated Pearson’s correlations (r) between MEPs amplitude and HbO responses (HbR, respectively). Furthermore, as cortical excitability is usually estimated on 20 or fewer MEPs while we had 75 measures, we applied a procedure to keep within-subject variability, that would have been lost by collapsing all trials in a single average. We then performed a bootstrap so that 2000 correlations were computed considering averaged amplitudes of 20 MEPs, 20 HbO and 20 HbR trials randomly selected for each run. The resulting r empirical distribution allowed the estimation of an average r-value and a confidence interval.

To estimate the effect of PAS on excitability (MEPs), HbO and HbR responses, we considered post-/pre- ratios. The effects of PAS across interventions were tested with a one-way ANOVA applied independently for MEPs, HbO and HbR ratios, whereas the effect of each intervention was tested with a one-sample t-test against 1.

To estimate the relationship between PAS-related excitability and hemodynamic changes, linear regressions were performed between MEP and HbO ratios (HbR, respectively). The regression was conducted on three classes: 1) pooling all sessions; 2) sessions exhibiting concordant effects only, defined as MEP and HbO and HbR ratios simultaneously larger or smaller than 1; and 3) sessions with discordant effects.

### 2.6 Data and code availability

The original raw data supporting the findings of this study are available upon reasonable request to the corresponding authors. fNIRS and TMS data were processed via Brainstorm software [Tadel et al., 2011] available at https://neuroimage.usc.edu/brainstorm/ and the fNIRS processing plugin - NIRSTORM (https://github.com/Nirstorm/nirstorm) in Brainstorm.

## 3. Results

### 3.1 Correlation between cortical excitability and task-related hemodynamic responses

Fig.3 illustrates the relationship between excitability (MEPs amplitude) and task-related hemodynamic activity (HbO and HbR). We found a significant positive linear correlation between MEP and HbO amplitude (r = 0.25, p=0.03) and a non-significant negative linear relationship between MEP amplitude and HbR (r = -0.16, p=0.171), meaning that an increased level of excitability corresponded to higher task-related HbO concentration and lower HbR concentration. These relationships were robust when considering the variability of the three amplitudes, as demonstrated by the bootstrapped correlation histograms (Fig.3c and d). The 95% confident interval of correlation value was [0.16, 0.33] for HbO and [-0.24, -0.08] for HbR. In both cases, the confidence interval did not cross the zero line, meaning that the variability of excitability and hemodynamic activity measures within subjects and across trials did not influence the sign of the correlation.

**Fig.3.**
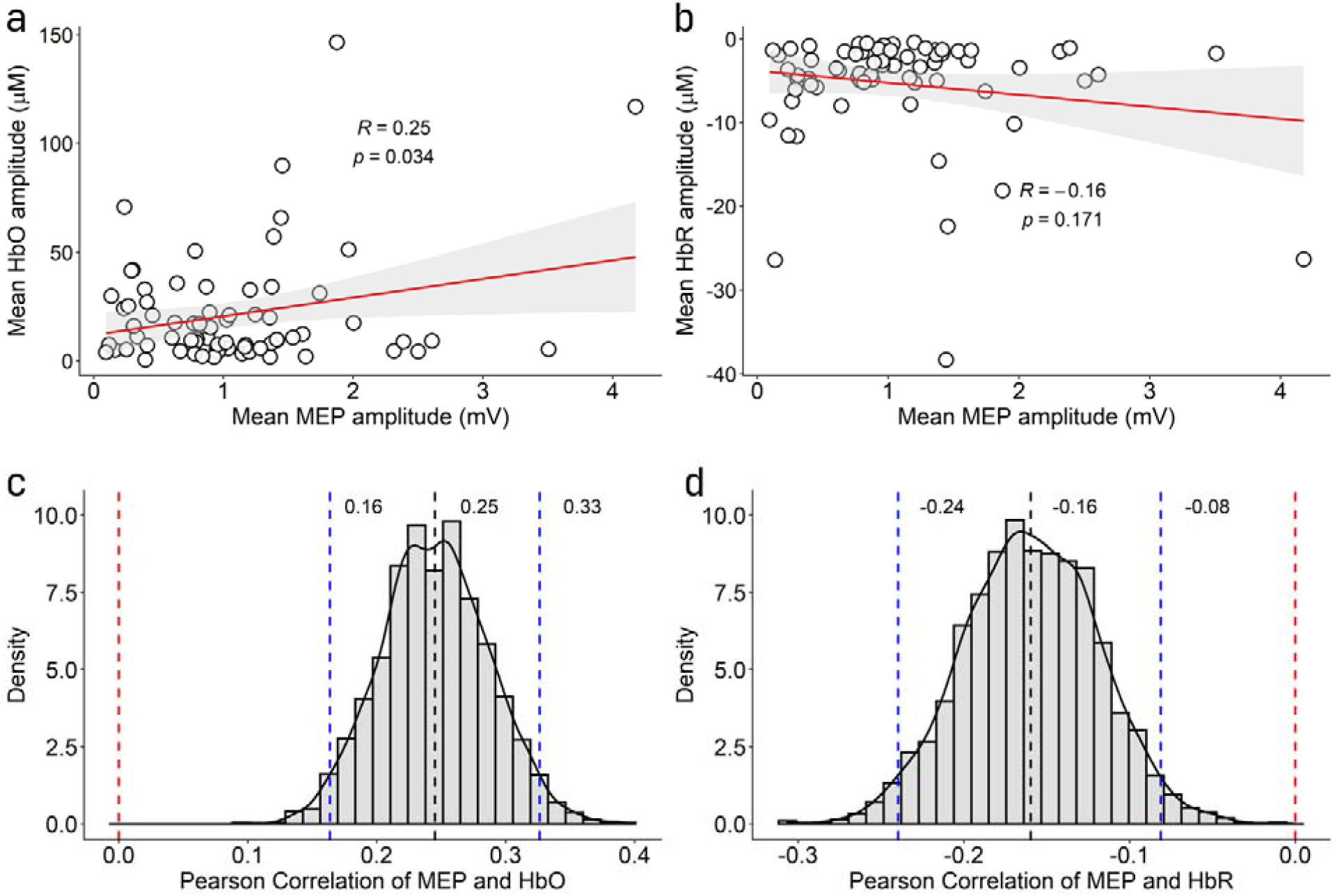
Correlation between MEP peak-peak amplitude and task-related HbO/HbR amplitudes. (a) scatterplot of HbO amplitude as a function of MEP amplitude. Each dot corresponds to a run (average over all MEPs amplitudes and all HbO trials). There was a significant positive linear relationship (Pearson’s r = 0.25, p=0.03). Red lines represent the estimated regression line, and the gray area indicated the 95% confidence interval. (b) There was a negative linear relationship between HbR and MEPs amplitude which did not reach statistical significance when considering run averages of all MEPs and HbR amplitudes. (c) and (d) the histogram of the correlation between MEP and HbO/HbR amplitudes, respectively, estimated from the bootstrap procedure (selecting 20 out of 75 MEP, HbO and HbR trials). Black curves represented the estimated density functions; black dashed lines showed the resulting mean correlation value; blue dashed lines indicated the 95% confidence interval estimated from the histograms; red dashed lines indicated r=0.

### 3.2 PAS effects on cortical excitability and task-related hemodynamic activity

Fig.4 shows the group-level MEP, HbO and HbR ratios for the PAS25, PAS10 and sham sessions. Overall, PAS produced similar effects for excitability and hemodynamic, with an overall increase of MEP, HbO and HbR ratios after PAS25, a decrease of MEP and HbO ratios after PAS10, and the ratio of sham was always in between PAS25 and PAS10. Table.1 summarizes the corresponding ratio values. Of note, all measures and especially fNIRS measures showed rather high variabilities, which likely prevented from reaching statistical significance (p>0.05 for the ANOVA and one-sample t-test) for the planned comparisons.

**Fig.4.**
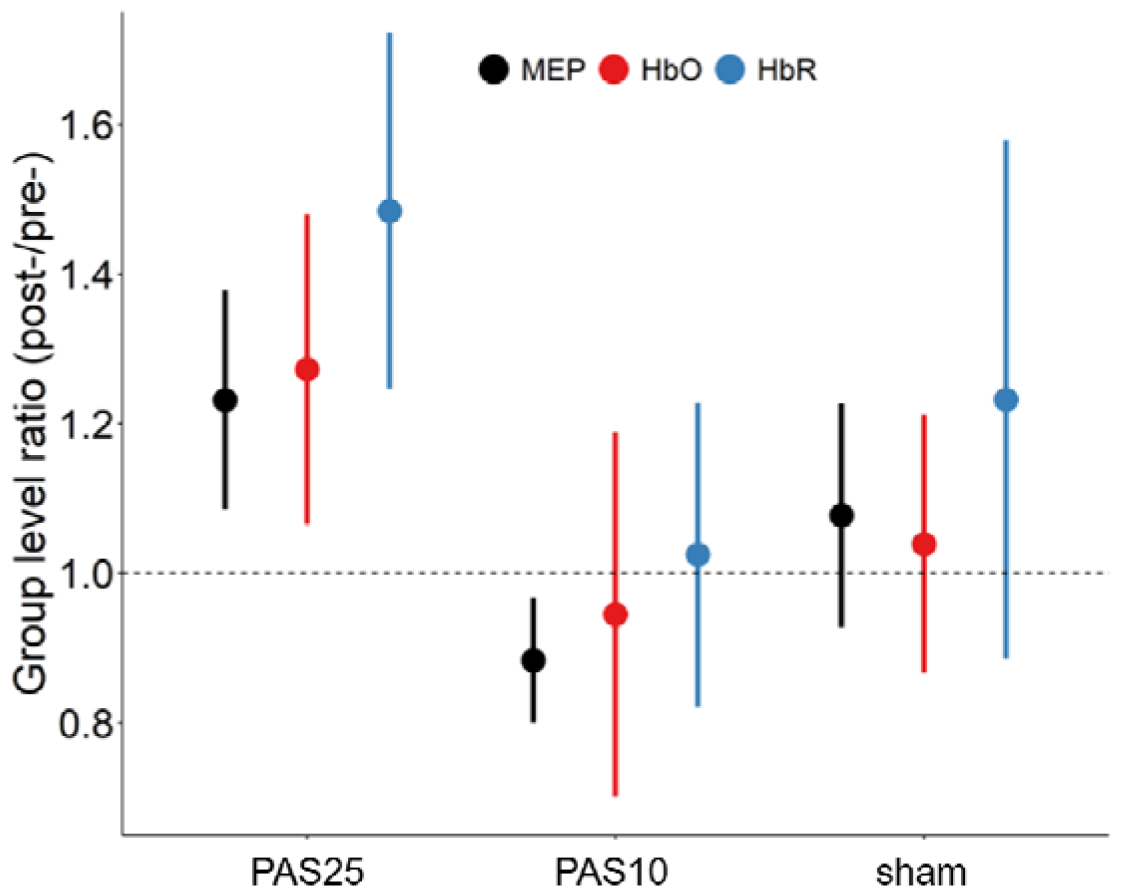
Group-level PAS effects on cortical excitability and hemodynamic activity. Effects of PAS25, PAS10 and sham on M1 cortical excitability (MEP amplitude) and M1 task-related hemodynamic activity (HbO and HbR) expressed as mean±SEM (standard error of the mean). PAS produced similar effects for excitability and hemodynamic activity, with an increase of MEP amplitude, HbO and HbR following PAS25, and a decrease of MEP amplitude and HbO following PAS10, a slight increase of the three measures after sham. All measures, and especially fNIRS measures, showed rather high variabilities (see standard error of the mean in the figure).

**Table.1.** Group-level PAS effects on cortical excitability and hemodynamic activity.

| | Ratio (Mean±SEM) | | |
| --- | --- | --- | --- |
|  | PAS25 | PAS10 | Sham |
| MEP | 1.23 ± 0.15 | 0.88 ± 0.08 | 1.08 ± 0.15 |
| HbO | 1.27 ± 0.21 | 0.95 ± 0.24 | 1.04 ± 0.17 |
| HbR | 1.48 ± 0.24 | 1.02 ± 0.20 | 1.23 ± 0.35 |

Fig.5 illustrates the single subject level (Sub02) PAS effects on reconstructed HbO and HbR. Following PAS25 and PAS10, absolute HbO and HbR amplitude were exhibiting increases and decreases, respectively. These effects were concordant with MEP amplitude changes (detailed values in Fig.5 caption). Fig.6 presented the group-level reconstruction maps. HbO peak amplitude (mean within the ROI,) increased after PAS25 and decreased after PAS10. HbR peak amplitude within the ROI decreased after PAS25 and also decreased after PAS10 (detailed values in Fig.6).

**Fig.5.**
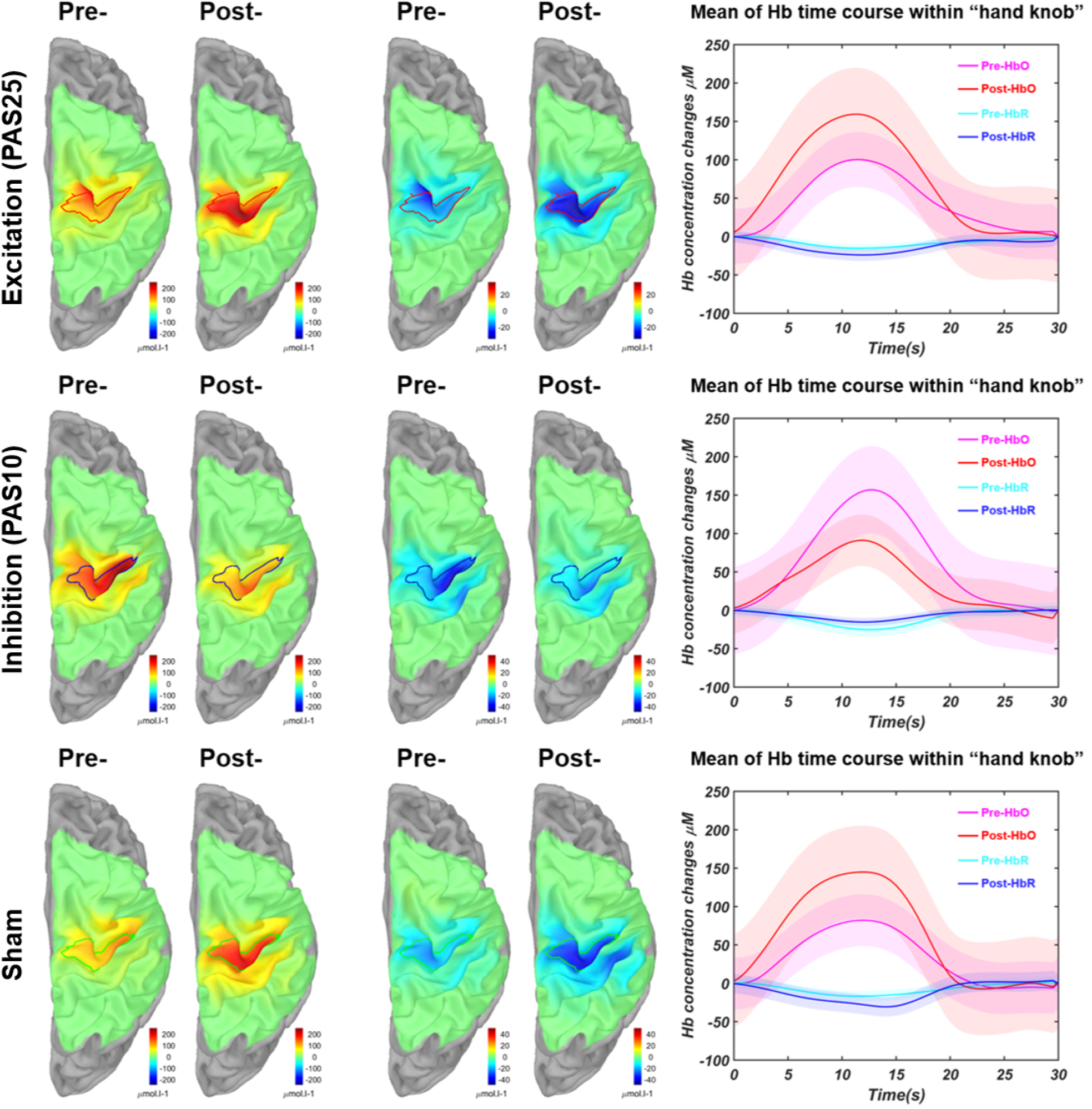
PAS effects on task-related hemodynamic responses using personalized fNIRS tomography for Sub02. Individual-level reconstructed fNIRS maps of HbO and HbR responses for each experimental session, pre- or post-intervention, PAS25 (1^st^ row), PAS10 (2^nd^ row), sham (3^rd^ row). Each map demonstrated the spatial distribution pattern of the task-related hemodynamic responses at its own peak timing of the reconstructed time courses showed in the last column. The red, blue and green profiles represented the extracted specific ROIs exhibiting significant hemodynamic responses at the peak amplitudes (see method section for details) for PAS25, PAS10 and sham, respectively. The averaged time course of HbO and HbR within each selected ROI for each session is presented in the last column by the solid lines, within a time window from 0s (the task onset) to 30s. All HbO and HbR time courses demonstrated the typical hemodynamic responses evoked by a 10s duration task. The shaded area represented the standard deviations of the time course within the ROI. Both the absolute amplitudes of HbO and HbR showed expected increase (after PAS25) and decrease (after PAS10) patterns within the selected ROIs. However, sham session also resulted in increased HbO and decreased HbR. The corresponding ratios of HbO and HbR were also in agreement with the mean MEP ratios, e.g., MEP ratio = 1.37, HbO ratio = 1.62 and HbR ratio = 1.56 for PAS25; then 0.85, 0.58 and 0.61 for PAS10, respectively. Colour maps for pre- and post- maps were fixed for specific hemoglobin and session (e.g., HbO in PAS25). HbO amplitudes generally exhibit a larger range than HbR amplitudes.

**Fig.6.**
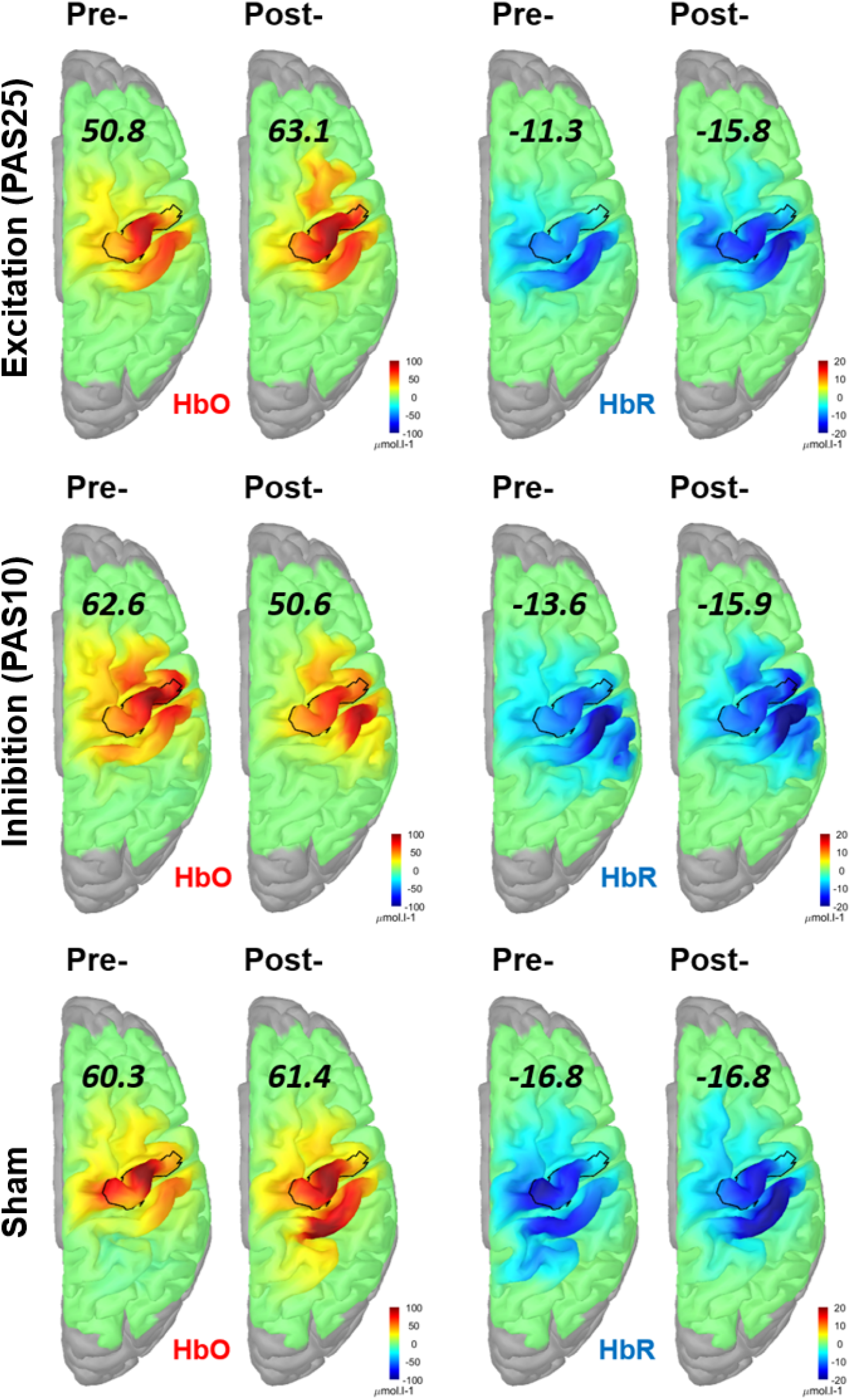
Group-level PAS effects on task-related hemodynamic responses. Group level reconstructed maps of HbO and HbR of each experimental session, pre- or post-, were shown in each row for PAS25, PAS10 and sham, respectively. Each map demonstrated the averaged corresponding individual-level task-related hemodynamic responses. Individual HbO/HbR maps selected at their respective peak were first coregistered on the mid-surface of the MNI ICBM152 template using Freesurfer spherical registration and then averaged over all subjects. The numbers indicated on every map are reporting the mean HbO or HbR amplitude within the corresponding ROI. This group-level ROI was defined manually along the M1 cortex of the template cortical surface to cover the hand knob region. Note that these maps were mainly considered for the visualization of PAS effects, whereas the statistical summary of hemodynamic changes was extracted at the individual levels. Colour maps for pre- and post- maps were fixed for specific hemoglobin and session (e.g., HbO in PAS25). HbO amplitudes generally exhibit a larger range than HbR amplitudes.

### 3.3 Relationship between PAS-related excitability and hemodynamic changes

When pooling all sessions together (Fig.7a), the resulted linear regression between HbO ratio (HbR, respectively) and MEP ratio presented a positive relationship, but no significant association (MEP-HbO ratios, r=0.19, p=0.29; MEP-HbR ratios, r=0.18, p=0.30). Fig.7b illustrated the linear regression for sessions with concordant PAS effects (e.g., MEP and HbO and HbR ratios were simultaneously larger or smaller than 1), resulting in significant positive linear corrections between MEP ratio and both HbO (r = 0.82, p<0.001) and HbR (r=0.88, p<0.001) ratios. No significant linear correlations were found when considering sessions with non-concordant PAS effects (Fig.7c).

**Fig.7.**
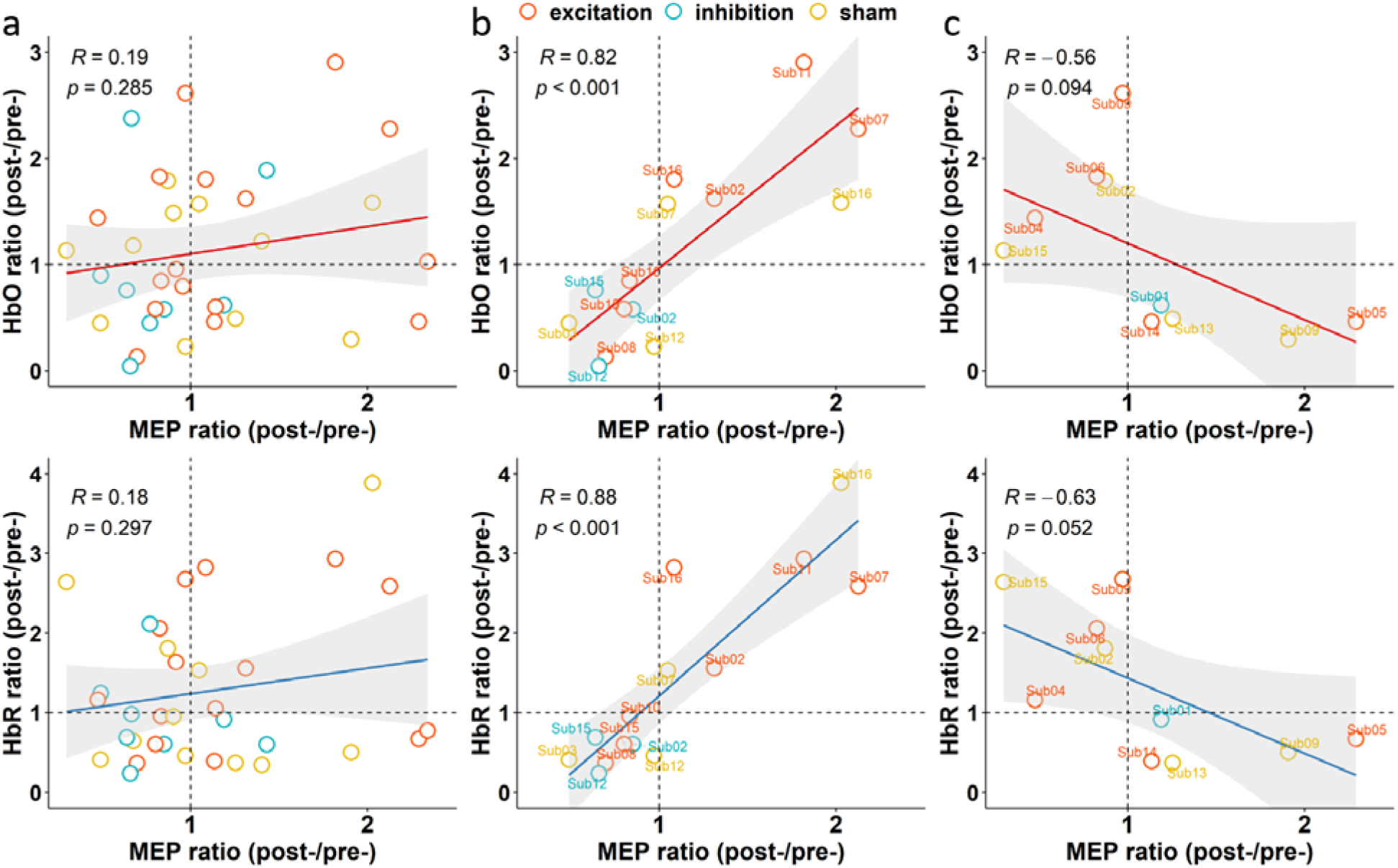
Correlates of task-related hemodynamic changes and PAS modulated excitability changes. The 1^st^ row presented HbO ratio as a function of MEP ratio and 2^nd^ row showed HbR ratio as a function of MEP ratio. Each point represented a session (e.g., PAS25 for Sub01, post-/pre- ratios). Sessions were color coded as red for PAS25, blue for PAS10 and yellow for sham. (a) linear regression between MEP ratios and HbO/HbR ratios when considering all sessions. We found positive Pearson’s correlations, r=0.19 and r=0.18 between MEP-HbO and MEP-HbR ratios, respectively, but none of them were statistically significant. (b) Linear regression for the PAS concordant sessions, in which MEP and HbO and HbR ratios were all larger than 1, or all smaller than 1. Both MEP-HbO and MEP-HbR cases demonstrated significant positive linear correlations (r=0.82 and r=0.88, p<0.001 in both cases). Besides, both fitted lines were not far from the point (1,1), e.g., HbO ratio = 0.96 and HbR ratio = 1.16, when fixing MEP ratio at 1, which is consistent with our prior knowledge. (c) Linear regression for the PAS non-concordant cases acting as a control in which none of the correlations were statistically significant. Two lines were also slightly more distant from the point (1,1), e.g., HbO ratio = 1.18 and HbR ratio = 1.43, when fixing MEP ratio at 1.

## 4. Discussion

Our study is the first one investigating the relationship between excitability and hemodynamic activity with simultaneous PAS-fNIRS in humans. We took advantage of this novel approach to achieve two main results: 1) fluctuations of cortical excitability were positively correlated with fluctuations of hemodynamic responses to the task; 2) there was a trend suggesting a positive linear relationship between effects of PAS on excitability and hemodynamic activity. Our results provide a unified view on two fundamental properties of cortical function. In addition, the demonstration of the effects of PAS on hemodynamic activity is relevant for the application of non-invasive brain stimulation techniques for the treatment of neuropsychiatric disorders. Finally, the tight link between excitability and hemodynamic activity may suggest that the effects on hemodynamics might be monitored via the standard spTMS technique.

### 4.1 Correlation between cortical excitability and hemodynamic activity

We demonstrated a link between excitability and task-related hemodynamic activity. Since PAS is known to induce variable effects across subjects, we pooled together all runs (before and after interventions) to investigate relationships between excitability and hemodynamic activity independently from specific PAS effects. Finger tapping is known to increase the metabolic demand and is therefore associated to increase HbO, decrease HbR and increase blood volume [Kashou et al., 2016; Novi et al., 2020]. Moreover, finger tapping itself is known to increase cortical excitability [Koeneke et al., 2006]. The correlation that we found underlines that metabolic demands linked to finger tapping depend on the excitability state when the task was performed. In other words, the metabolic demands seem state-dependent, where brain state corresponded here to cortical excitability. State dependency is a very well-known concept in cortical function and involves multiple measures of neuronal activity such as activity, oscillations and connectivity [Giambattistelli et al., 2014; Gonçalves et al., 2006; Romei et al., 2008; Silvanto et al., 2008; Silvanto and Pascual-Leone, 2008].

### 4.2 PAS effects on hemodynamic activity

In the field of non-invasive human brain stimulation, two previous studies investigated the effect of M1 cortical excitability modulation on task-related hemodynamic activity. Kriváneková et al (2013) combined PAS and “offline” fMRI and reported no definite effects of PAS on either the task-related BOLD signal of the sensorimotor regions or resting-state functional connectivity. They reported that BOLD fluctuations following PAS were rather unpredictable, with almost no change after excitatory PAS, BOLD increase after inhibitory PAS and BOLD decrease after sham. Comparing these results to ours is challenging, but the low sampling frequency of fMRI might have contributed to their negative results. A low sampling rate means limited temporal sampling which might dilute small effects of excitability changes that we identified mostly around the peak of the hemodynamic response. Although PAS effects on HbO and HbR were also not significant in our study, we did find expected trends in some individuals and at the group level (Fig.4, 5 and 6). Our results are certainly in agreement with those reported by Chiang et al., (2007), who combined rTMS and fNIRS. They found that 1Hz rTMS over M1 induced an expected HbO increase in the contralateral cortex lasting up to 40 minutes, likely related to reciprocal inhibition mechanisms [Di Lazzaro et al., 2014]. Such cortical excitability effects on hemodynamic were also observed in other cortical regions. 5Hz rTMS on the right parietal cortex during the retention period of a match-to-sample task significantly increased HbO levels during the task period [Yamanaka et al., 2010]; whereas continuous theta-burst stimulation (cTBS) applied on the left dorsolateral prefrontal cortex (DLPFC) reduced Emotional Stroop task-evoked HbO levels bilaterally [Tupak et al., 2013]; cTBS on the right-DLPFC also reduced HbO during the dictator game [Maier et al., 2018].

Further development of our simultaneous fNIRS-TMS protocol may open new avenues for understanding the mechanism of PAS per se. For instance, the sparse rhythmic stimulation involved during PAS will allow investigating pulses-related hemodynamic effects and their build-up. Such analysis was out of our scope and will be considered in future investigations.

### 4.3 Correlation between PAS effects on excitability and PAS effects on hemodynamic activity

We also evaluated the correlation between excitability changes and hemodynamic activity changes modulated by interventions. Such analysis was challenging as the modeling involved dealing with the variabilities of MEP amplitudes, hemodynamic activity and PAS effects. Similar to other neuromodulatory techniques, PAS is known for inducing variable and sometimes unpredictable or even reversed effects [Suppa et al., 2017; Ziemann and Siebner, 2015], depending upon multiple factors, including state of the brain, genetic susceptibility, position of the coil, and much more [Ziemann and Siebner, 2015]. Using fMRI and PAS, Kriváneková et al., (2013) did not report any significant relationship between PAS related excitability changes and finger-tapping related BOLD changes. We found similar results regarding the correlation between MEP ratios and HbO (HbR, respectively) ratios when considering all sessions. Note that in our first correlation analysis we found that fluctuations of cortical excitability were positively correlated with fluctuations of hemodynamic responses to the task. Then if the variability of MEP amplitudes, hemodynamic activity and PAS effects do not introduce variability on ratio calculations of MEP and HbO/HbR, similar significant positive correlations should have been observed. Hence, we believe that these variabilities deflated the correlation value in our second correlation analysis. To further address this point, we constrained this correlation analysis for sessions with concordant (Fig.7b) or discordant (Fig.7c) PAS effects on MEP, HbO and HbR ratios. This proposed strategy simply regulates the influences of data variability on the correlation of ratios by claiming that the correlation should have existed in sessions where those three measurements varied in the same direction after PAS (i.e., less data variability). Interestingly, there was a significant correlation between MEP ratios and HbO/HbR ratios. As a contrast, sessions with discordant PAS effects (Fig.7.c) did not show significant correlations. Moreover, when data exhibit less variability, HbO or HbR ratio should be close to 1 when fixing MEP ratio at 1. In concordant case (Fig.7b), when fixing MEP ratio at 1 the fitted lines indicate that HbO ratio (0.96) and HbR ratio (1.16) were actually closer to 1 than in discordant case (HbO ratio = 1.18 and HbR ratio = 1.43, Fig.7c). Despite the simplicity of this regulation strategy using concordance, these evidences suggest that further advanced data analysis which better handles variability in the data may reveal a solid correlation between PAS effects on MEP and its effects on HbO/HbR. To summarize, our finding on this second correlation analysis underlines the complexity of non-invasive brain stimulation effects, which are often only investigated in the domain of excitability, but always involve also hemodynamic activity, electromagnetic activity, connectivity, and much more [Pellegrino et al., 2018; Pellegrino et al., 2019]. This finding also underlines the tight link between these cortical properties and may offer new opportunities for patients’ treatment whenever the target action is a modulation of blood and hemoglobin supply.

### 4.4 Reliability and robustness

We did our best to acquire and analyze data in a robust way, including only male subjects, tuning PAS on individual N20, applying neuronavigation on individual MRI, collecting many MEPs (75 vs usual 20 trials). We considered personalized fNIRS data acquisition targeting the hand-knob using an optimal montage [Machado et al., 2014; Machado et al., 2018; Pellegrino et al., 2016] computed on the individual MRI. Physiological noise such as heartbeats, respiration and Mayer wave were minimized by filtering and regression using short-distance channels that only probe hemodynamics in the scalp [Zeff et al., 2007]. Optodes were positioned with neuronavigation and glued on head skin via collodion to ensure good contact and optimal probe design. We extracted HbO/HbR features after 3D reconstructions along the cortical surface. fNIRS 3D reconstructions have been shown to provide more accurate quantification [Arridge, 1999; Boas et al., 2001] of HbO/HbR than sensor level analyses applied in most fNIRS involved TMS studies [Curtin et al., 2019b; Curtin et al., 2019a; Oliviero et al., 1999; Thomson et al., 2011; Thomson et al., 2013]. More importantly, rigorous statistical procedures were conducted considering variabilities of data at different levels: 1) a bootstrap procedure to pool together data from all recordings instead of simply taking averages to investigate the relationship between excitability and hemodynamic activity; and 2) a resampling technique ensured to extract reliable, robust and data driven (intervention type blind) HbO/HbR measures from fNIRS reconstructions. Finally, most of fNIRS involved TMS studies only reported results on HbO. However, we showed consensus results when considering both HbO and HbR signals, as recommended in a recent fNIRS guideline paper [Yücel et al., 2021].

### 4.5 Limitations

The small number of subjects and unbalanced data set are the main limitations of this study. However, considering the main contribution of this study was investigating the correlation, the number of sessions involved in the correlation analysis (Fig.3 and Fig.7) was sufficient. Secondly, the finger-tapping task and the spTMS session were not conducted at the same time, but within a few minutes. To be noted, applying TMS during the task would have answered different biological questions than the ones assessed here. Nonetheless, the time gap between TMS and motor tasks may have introduced some noise when investigating the correlation between MEP and HbO/HbR because of the fluctuations of both measures over time. Finally, more advanced data analysis which better handles data variability may help to reveal a significant correlation between PAS effects on M1 excitability and its effects on task-related hemodynamic activity.

## 5. Conclusions

In conclusion, we demonstrated a linear relationship between brain excitability and task-related hemodynamic activity measured using personalized fNIRS. We also demonstrated that PAS may have effects on hemodynamic activity in addition to those on excitability and also influences the relationship between excitability and activity. These effects are not necessarily PAS-specific and may characterize other non-invasive brain stimulation techniques as well. Finally, our findings may further expand the field of non-invasive brain stimulation application for treating brain disorders by targeting those areas for which a modulation of hemodynamic activity is desired.

## Acknowledgments

This work was supported by the Natural Sciences and Engineering Research Council of Canada Discovery Grant Program (CG and JML), an operating grant from the Canadian Institutes for Health Research (CIHR MOP 133619 (CG)), a FRQNT research team grant and a FRQS-Quebec Bio-Imaging Network (QBIN) Pilot Project grant. fNIRS equipment was acquired using grants from NSERC Research Tools and Instrumentation Program and the Canadian Foundation for Innovation (CG). SF is supported by the Canada Research Chair in Cognitive Neuroplasticity. ZC is funded by the Fonds de recherche du Québec – Sante (FRQS) Doctoral Training Scholarship and the PERFORM Graduate Scholarship in Preventive Health Research. GP is funded by Strauss Canada Foundation and a McGill/MNI - Fred Andermann EEG and Epilepsy Fellowship. MU is funded by the Horizon Postdoctoral Fellowships of Concordia University.

## Conflict of interest

The authors declare no potential conflict of interest.

## Author contributions

**ZC:** Formal analysis, Methodology, Data collection, Original draft preparation; Reviewing and Editing **GP**: Conceptualization, Investigation, Original draft preparation; Reviewing and Editing **AS, ED, MU and CA:** Data collection, Reviewing and Editing; **JML:** Methodology, Reviewing and Editing; **SF:** Conceptualization, Reviewing and Editing; **CG:** Conceptualization, Methodology, Supervision, Reviewing and Editing.

## Appendices

## A: Experiment design and Data acquisition

### Anatomical MRI

To guide TMS procedure and to calculate the head model for fNIRS acquisitions and analyses, MR brain images were acquired for each participant, using a General Electric Discovery MR750 3T scanner at the PERFORM Center of Concordia University, Montréal, Canada. T1-weighted images were acquired using the 3D BRAVO sequence and the following parameters: 1 × 1 × 1 mm^3^, 192 axial slices, 256 × 256 matrix. T2-weighted anatomical images were acquired using the 3D Cube T2 sequence and the following parameters: 1 × 1 × 1 mm^3^ voxels, 168 sagittal slices, 256 × 256 matrix.

### fNIRS data acquisition

fNIRS data were acquired using a Brainsight fNIRS machine (Rogue-Research Inc, Montréal, Canada) sampling at 10Hz. A neuro-navigation system (Rogue-Research Inc, Montréal) with individual MRI images guided the installation of the optodes according to the previously estimated optimal montage (see Fig.2). Optodes were glued over the subject’s scalp with collodion, taking care to remove the hair between skin and sensor, thus allowing reducing motion artifacts during prolonged recordings (Fig.2) [Machado et al., 2018; Pellegrino et al., 2016; Yücel et al., 2014].

### Measurement of M1 cortical excitability

TMS was delivered with a Magstim 200^2^ stimulator (Magstim Company, Carmarthenshire, Wales, UK) connected to a figure-8 coil (Magstim double 70mm remote control coil). Subjects were sitting in a comfortable armchair with the support of the left arm and neck (see Fig.2.d). To reduce potential motion, we placed the head of the participant between a mechanical arm wrapped with a soft cushion and the TMS coil (see Fig.2.d). TMS procedures were guided by the neuro-navigation system Brainsight (Rogue-Research Inc, Montréal, Canada). TMS coil was placed tangential to the scalp and with a 45° angle to the midline of the head [Thomson et al., 2013]. MEPs were recorded by a BrainAmp ExG bipolar system with 2 TECA disposable 20mm disk electromyography (EMG) electrodes attached on the right abductor pollicis brevis (APB), with a standard belly-tendon montage (Fig2.d). The TMS “hot spot” was found for each session as the location with the maximal Motor Evoked Potentials (MEP). Resting motor threshold (RMT) was defined efficiently using the TMS Motor Threshold Assessment Tool (MTAT 2.0, http://www.clinicalresearcher.org/software.htm) based on maximum-likelihood parameter estimation by sequential testing approach [Ah Sen et al., 2017; Awiszus et al., 1999].

### Somatosensory evoked potentials

Electrical stimulation (Digitimer DS7A, Letchworth Garden City, U.K) at the left wrist (e.g., Median Nerve) was performed after the MRI scan. Two bipolar EEG (BrainAmp ExG, Brain Products GmbH, Germany) electrodes located at CP3 and CP4 were used to measure subject specific N20 latency. Stimulation was delivered slightly above the motor threshold at 4Hz for 2 minutes. N20 latency was visually estimated via the online segment averaging functionality of the BrainVision Recorder (BrainAmp ExG, Brain Products GmbH, Germany).

## B: Reliable and robust estimation of task-related hemodynamic responses using fNIRS 3D reconstructions and resampling technique

To conduct a reliable and robust task-related hemodynamic response estimation to compare hemodynamics before and after interventions, we combined fNIRS 3D reconstruction and resampling techniques to appropriately handle variability between trials and influence of eventual motion artifacts. The workflow consisted of 3 steps: a) resampling of “all the possible” averaged optical density time courses; b) fNIRS 3D reconstruction along the cortex using the MEM and c) definition of a session specific spatial ROI for HbO/HbR features extraction.

a. We conducted a resampling of “all the possible” averaged optical density time course (−10s to 30s) as the input for further fNIRS 3D reconstructions. First, for every 20 blocks of pre-processed optical density data, we sub-averaged all 16 out of 20 blocks, resulting in a total of 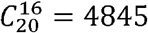 possible combinations. These averaged trials were then sorted by the averaged signal-to-noise ratio (SNR) for two wavelengths. SNR was calculated by the largest amplitude among all channels from 0s to 30s, normalized by the mean of standard deviation over all channels during the baseline [-10s, 0s]. Finally, we selected 101 out of 4845 sub-averaged blocks centred around the median SNR of all sub-averages. The selection of 16 trials for sub-averaging was empirically defined according to the observation that usually there were around 4 artifacts contaminated blocks per condition (i.e., containing eventual motion artifacts). For artifacts contaminated data, large motion artifacts would then result in large SNR of sub-averaged trials. On the other hand, for data that are not contaminated by artifacts, the SNR distribution will be flat. Therefore, selecting sub-averaged trials around the median SNR should ensure a good representation of fNIRS responses, discarding artifact sub-averages. We finally chose 101 sub-averaged trials to ensure a good representation of the underlying distribution of SNR values, while being sensitive to the inherent variability of task-evoked fNIRS responses.
b. fNIRS 3D reconstruction along the cortex was conducted using the MEM method applied to each of 101 sub-averaged trials specific for task run (e.g., pre-PAS25 of Sub01). We, therefore, took into account the variability of the hemodynamic response, instead of considering only one averaged response. “All the possible” spatiotemporal maps of HbO and HbR along the cortical surface were reconstructed using MEM from these 101 resampled sub-averaged trials. MEM is an efficient nonlinear probabilistic Bayesian framework to incorporate prior knowledge in the solution of the inverse problem for 3D reconstruction [Amblard et al., 2004]. We have demonstrated the excellent accuracy of MEM, especially its high sensitivity and specificity to the spatial extent of the underlying generators in the context of electro-/magneto-encephalogram source imaging [Chowdhury et al., 2013; Chowdhury et al., 2016; Grova et al., 2016; Hedrich et al., 2017; Heers et al., 2016; Pellegrino et al., 2020] as well as for fNIRS 3D reconstructions [Cai et al., 2021].
c. A session specific (e.g., PAS25 of Sub01) spatial ROI was finally defined along the cortical surface, to extract the reconstructed HbO/HbR time courses features from MEM reconstructed spatiotemporal maps. To do so, we first extracted the task run specific (e.g., pre-PAS25 of Sub01) HbO/HbR peak maps, at the peak timing estimated from the reconstructed time courses within the hand knob. This resulted in 101 HbO/HbR peak spatial maps for each task run. Then, a one-sample t-test of HbO (respectively HbR) amplitude for each vertex of the map was performed across 101 maps for each task run. Regions exhibiting a significant response (p<0.05, false discovery rate corrected) were kept as the cortical area, which contained significant hemodynamic responses of one task run. For each session (e.g., PAS25 of Sub01), this analysis resulted in 4 regions exhibiting significant hemodynamic responses, i.e., HbO and HbR response t-maps, in pre- and post-intervention conditions. The final ROI was defined as the intersection between these 4 statistically significant regions to ensure reliability and robustness. Note that this final ROI was confirmed to be within the M1 cortical area for all participants. Two PAS10 sessions and 1 sham session were rejected due to the empty intersection regions (no overlapping between the 4 regions), suggesting a lack of reliability of the resulting hemodynamic responses. We defined this ROI for each intervention session rather than for each subject, considering the variability of task performance within each subject, since the intervention sessions of each subject were performed on different days.

## C: PAS effects on cortical excitability and hemodynamic activity

As shown in Fig.C1, PAS exerted the expected group level effect with average MEP ratios (mean±SEM) of 1.23±0.15, 0.88±0.08 and 1.08±0.15 for PAS25, PAS10 and Sham, respectively. Such group level trends were also manifested in HbO and HbR ratios. For instance, the group level effect with average HbO ratios (mean±SEM) were 1.27±0.21, 0.95±0.24 and 1.04±0.17 for PAS25, PAS10 and Sham, respectively; and the group level effect with average HbR ratios (mean±SEM) were 1.48±0.24, 1.02±0.20 and 1.23±0.35 for PAS25, PAS10 and Sham, respectively. However, these effects did not reach statistical significance (one-sample t-test against 1, p>0.05). There were also no significant differences among 3 interventions (ANOVA test p>0.05).

**Fig.C1.**
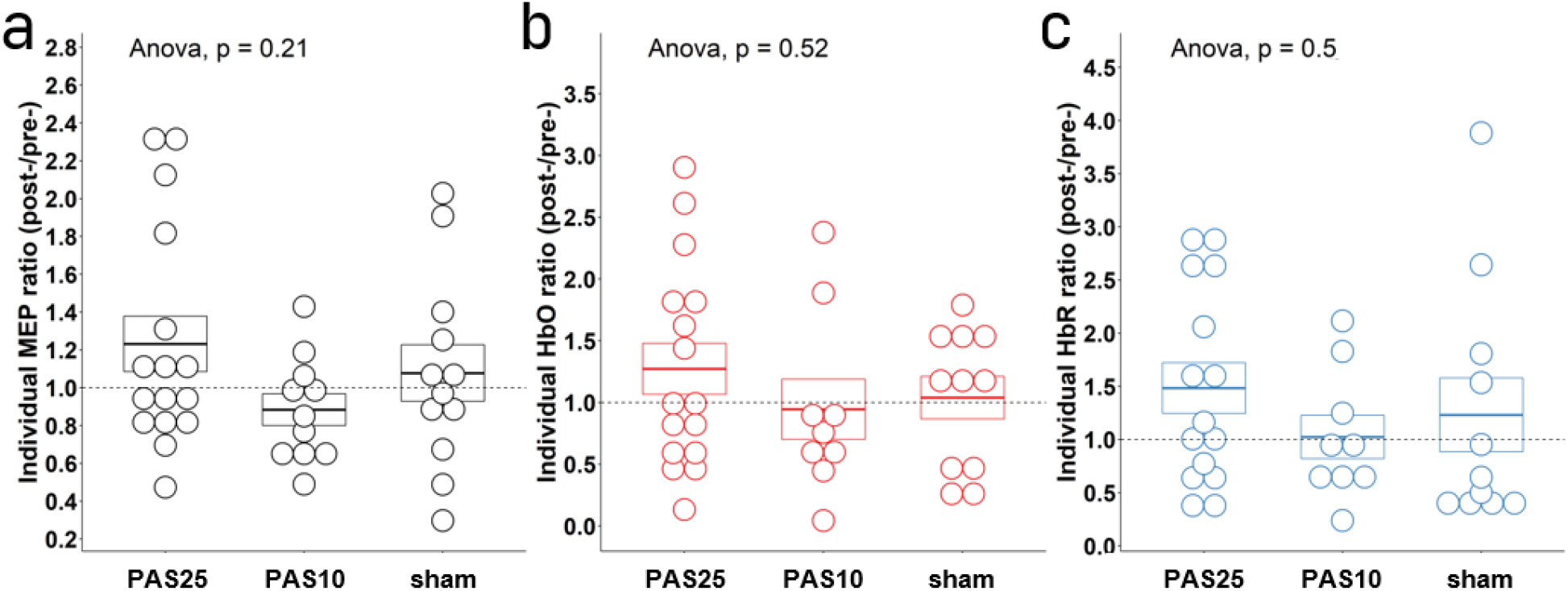
PAS effects on cortical excitability and hemodynamic activity. At the group level, Mean±SEM (standard error of the mean) for the excitatory, inhibitory and sham session were: (a) 1.23±0.15, 0.88±0.08 and 1.08±0.15 for MEP ratios; (b) 1.27±0.21, 0.95±0.24 and 1.04±0.17 for HbO ratios; and (c) 1.48±0.24, 1.02±0.20 and 1.23±0.35 for HbR ratios. Even if trends were observed, none of these ratios were significantly different from 1 based on a one-sample t-test. The individual ratios represented by each dot demonstrated a relatively large variance of the ratios for each scenario. Boxes showed the Mean±SEM in each case. Note that one MEP session (e.g., Sub16, PAS10) was rejected from the whole analysis due to a high ratio of 2.81; and one HbR session (e.g., Sub12, PAS25, HbR) was rejected from the whole analysis due to a high ratio of 5.45. A similar process was also considered in Kriváneková et al (2013).

## Notes

### Competing Interest Statement

The authors have declared no competing interest.

